# Chicken IRF10 suppresses the cGAS-STING-IFN antiviral signaling pathway by targeting IRF7

**DOI:** 10.1101/2025.11.05.686717

**Authors:** Sen Jiang, Jingyu He, Kun Wang, Mengjia Lv, Hongjian Han, Bingjie Sun, Nanhua Chen, Wanglong Zheng, Jianzhong Zhu

## Abstract

The chicken (ch) innate immune cGAS-STING-IFN signaling pathway plays a crucial role in host antiviral defense, wherein chIRF7 serves as the key downstream transcription factor of chSTING responsible for initiating type I IFN transcription. While chicken IRF family encompasses 8 members, the potential involvement of other IRF members in chicken cGAS-STING-IFN signaling transduction and/or regulation remains poorly characterized. In this study, we identified chIRF10 as a potent suppressor of chicken cGAS-STING-IFN signaling against several viruses, by screening of chIRF family members. Mechanistically, the inhibitory activity of chIRF10 depended on its IRF associated domain (IAD) but not its DNA binding domain (DBD). Further, chIRF10 suppressed both cGAS-STING mediated IFN signaling and TBK1/IKKε/IRF7 activated IFN signaling. Importantly, chIRF10 could interact with chIRF7, and inhibit its dimerization and activation. In summary, our findings unveiled a novel function of chIRF10 as well as a novel regulatory mechanism governing chicken cGAS-STING-IFN antiviral signaling, thereby providing a scientific foundation for developing strategies against chicken viral diseases.

## Introduction

Innate immunity serves as the first line of defense for the organism to recognize and resist pathogenic microorganism infections. Years of researches have revealed that innate immunity constitutes an extensive network system composed of multiple pattern recognition receptor (PRR) families, including Toll-like receptors (TLRs), C-type lectin receptors (CLRs), RIG-I-like receptors (RLRs), NOD-like receptors (NLRs), and cytosolic DNA sensors. When pathogen-associated molecular patterns (PAMPs) released by invading microorganisms or endogenous damage-associated molecular patterns (DAMPs) are recognized by PRR, the corresponding signaling pathways relay signals in a cascading manner, leading to the activation of type I interferons (IFNs), inflammatory cytokines, and chemokines, thereby initiating innate immune responses (1, 2).

DNA sensors include TLR9 and cytosolic DNA sensors (3). Cyclic GMP-AMP synthase (cGAS), identified in 2013 as a cytosolic DNA sensor, plays a critical role in recognizing cytosolic DNA (4). Current studies have also reported the nuclear localization of cGAS, where it performs multiple functions (5, 6). In the cytoplasm, cGAS detects viral double-stranded DNA (dsDNA) as well as aberrant endogenous DNA, such as nuclear and mitochondrial DNA (7). As a member of the nucleotidyltransferase family, cGAS catalyzes the synthesis of a unique cyclic dinucleotide, 2’3’-cGAMP, from ATP and GTP upon binding dsDNA. This second messenger rapidly diffuses throughout the cytoplasm and specifically binds to STING on the endoplasmic reticulum (ER), leading to its activation (8–10). Activated STING undergoes a conformational change that exposes the TBK1 and IRF3 recruitment sites within its C-terminal tail (CTT). Subsequently, TBK1 undergoes autophosphorylation and phosphorylates both STING and the recruited IRF3. Phosphorylated IRF3 forms homodimers, translocates into the nucleus, and initiates the transcription of type I interferons (IFN) (11). STING can also moderately activate nuclear factor kappa B (NF-κB) through the IKKα/β/γ complex, which cooperates with IRF3 to enhance IFN production and promote the expression of pro-inflammatory cytokines (12).

Furthermore, STING has been implicated in regulating cellular homeostasis, including processes such as autophagy, apoptosis, necroptosis, and programmed necrosis (13, 14). The chicken cGAS-STING signaling pathway also plays a vital role in defending against various viral infections (15, 16). Notably, due to the natural absence of IRF3 in chickens, chicken IRF7 substitutes for IRF3 in activating the transcription and expression of type I IFNs (17, 18).

Interferon regulatory factors (IRFs) were originally identified as regulators of type I IFNs and IFN-stimulated genes (ISGs). In fact, IRFs represent a class of multifunctional transcription factors that play important roles in antiviral responses, inflammatory reactions, as well as cell differentiation and development (19). To date, nine IRF family members have been identified in mammals, including IRF1 (MAR), IRF2, IRF3, IRF4 (LSIRF), IRF5, IRF6, IRF7, IRF8 (ICSBP), and IRF9 (ISGF3γ or p48), while IRF10 has been found in chickens (19, 20). All IRFs contain a highly conserved DNA-binding domain (DBD) at the N-termini. With the exception of IRF6, all other members also contain a relatively conserved IRF associated Domain (IAD) at the C-termini (21). Unlike other IRF members, IRF1 and IRF2 possess a unique IAD (22). The DBD enables IRFs to recognize and bind specific Interferon-Stimulated Response Element (ISRE) promoter sequences (A/GNGAAANNGAAACT) (23). The IAD is responsible for the formation of homodimers or heterodimers among IRFs or with other transcription factors, thereby facilitating transcriptional activation or regulation (24).

Members of the IRF family play crucial roles in the regulation of interferon (IFN) signaling and the coordination of immune responses. IRF1, the first identified member of the IRF family, is constitutively expressed and can be induced by IFN-γ (25). IRF2 was identified through cross-hybridization with IRF1 cDNA (26). IRF1 recognizes and binds to the interferon-sensitive response element (ISRE) promoter sequence, directly activating the transcription of type I IFNs and interferon-stimulated genes (ISGs) (27). IRF2 is also constitutively expressed but can be induced by viruses and IFNs (26). The N-terminal domain of IRF2 shares high homology with that of IRF1 and recognizes the same promoter DNA sequence. As a result, IRF2 acts as an antagonist of IRF1 by competing for binding to the promoter sequence, thereby suppressing the transcriptional activity of IRF1 (26). IRF3 serves as a key transcription factor in innate immune signaling pathways, such as cGAS-STING and RIG-I/MDA5–MAVS, where it activates IFN-β expression by binding to the IFN-β promoter DNA sequence and initiating its transcription (19). IRF7 shares high amino acid homology with IRF3 and can directly bind the ISRE promoter sequence, performing functions similar to those of IRF3, particularly in activating IFN-α transcription (28). IRF4 is the only IRF not regulated by IFN and is involved in cell development and differentiation (29). IRF8, a homolog of IRF4, is constitutively expressed and primarily participates in cell development, differentiation, and immune regulation (30–33). IRF5 and IRF6 are homologous proteins; IRF5 has been shown to be involved in the induction of IFN-α, while the function of IRF6 is unrelated to IFN signaling and is mainly associated with cellular development (19, 34). IRF9, also known as ISGF3γ, is expressed in a variety of cells. As a critical component of type I IFN signaling, it forms the transcription complex ISGF3 together with STAT1 and STAT2, which collectively activates the transcription of ISGs (35). Chicken (ch) IRF family members are highly conserved with their mammalian counterparts but exhibit species-specific differences (17, 36–38). As mentioned earlier, chickens lack IRF3, and its function is compensated by IRF7 (39). Consequently, chIRF7 is often targeted by various viruses to suppress innate immune responses and facilitate immune evasion (39–42). Moreover, chickens naturally lack IRF9 but possess a unique IRF10 (20, 39). Although some functions of chicken IRFs have been studied, many aspects remain poorly understood, and the roles of chIRFs other than chIRF7 require further investigation.

In this study, we investigated the relationship between chicken IRF family members and the chicken cGAS-STING-IFN signaling pathway. It was found that only chIRF7 mediates chicken cGAS-STING activated IFN transcription, which is consistent with our previous findings. Moreover, we discovered that chIRF10, a chicken-specific member of the IRF family, acts as a negative regulator by suppressing the cGAS-STING-IFN antiviral signaling pathway. Further results demonstrated that chIRF10 significantly inhibits chIRF7 mediated activation of antiviral IFN signaling by directly interacting with IRF7 via its IAD domain, thereby suppressing IRF7 activation.

## Results

### Screening of chicken IRF family members for involvement in the chicken cGAS-STING-IFN signaling pathway

Previous studies including ours revealed that chicken naturally lack IRF3, and instead utilize IRF7 as the key transcription factor downstream of STING to activate type I IFN transcription (17, 18). However, the chicken IRF family comprises multiple members, and whether other IRFs participate in or regulate the chicken cGAS-STING-IFN antiviral signaling pathway remains unclear. To investigate this, we constructed expression vectors for eight members of the chicken (ch) IRF family including IRF1, IRF2, IRF4, IRF5, IRF6, IRF7, IRF8 and IRF10. Since 293T cells lack basal IRF7 expression, and chicken cGAS-STING cannot activate IFN signaling in this cell line (17). Therefore, we first conducted experiments in 293T cells and found that, among all chicken IRFs, only IRF7 synergistically activated the ISRE promoter in combination with chicken cGAS-STING (Fig 1A). Secondly, in the chicken fibroblast cell line DF-1 with basal IRF7 expression (18), chIRF7 enhanced chSTING activated IFN-β promoter (Fig 1B), confirming the role of chIRF7 as a transcription factor mediating chSTING activated IFN induction (17, 18). Further, it was found that the chIRF10 significantly suppresses chSTING activated IFN-β promoter (Fig 1B). To validate these results, we repeated the experiment in DF-1 cells using pEGFP-C1 based IRFs other than p3×FLAG-CMV expressed IRFs, and consistent results were yielded (Fig 1C). Together, these screening generated results suggested that among the chIRFs tested, only chIRF7 mediates IFN transcription upon activation of chicken cGAS-STING, confirming previous finding. Notably, we identified the chIRF10 as a significant inhibitor of the IFN signaling activated by chicken cGAS-STING.

**Figure 1.**
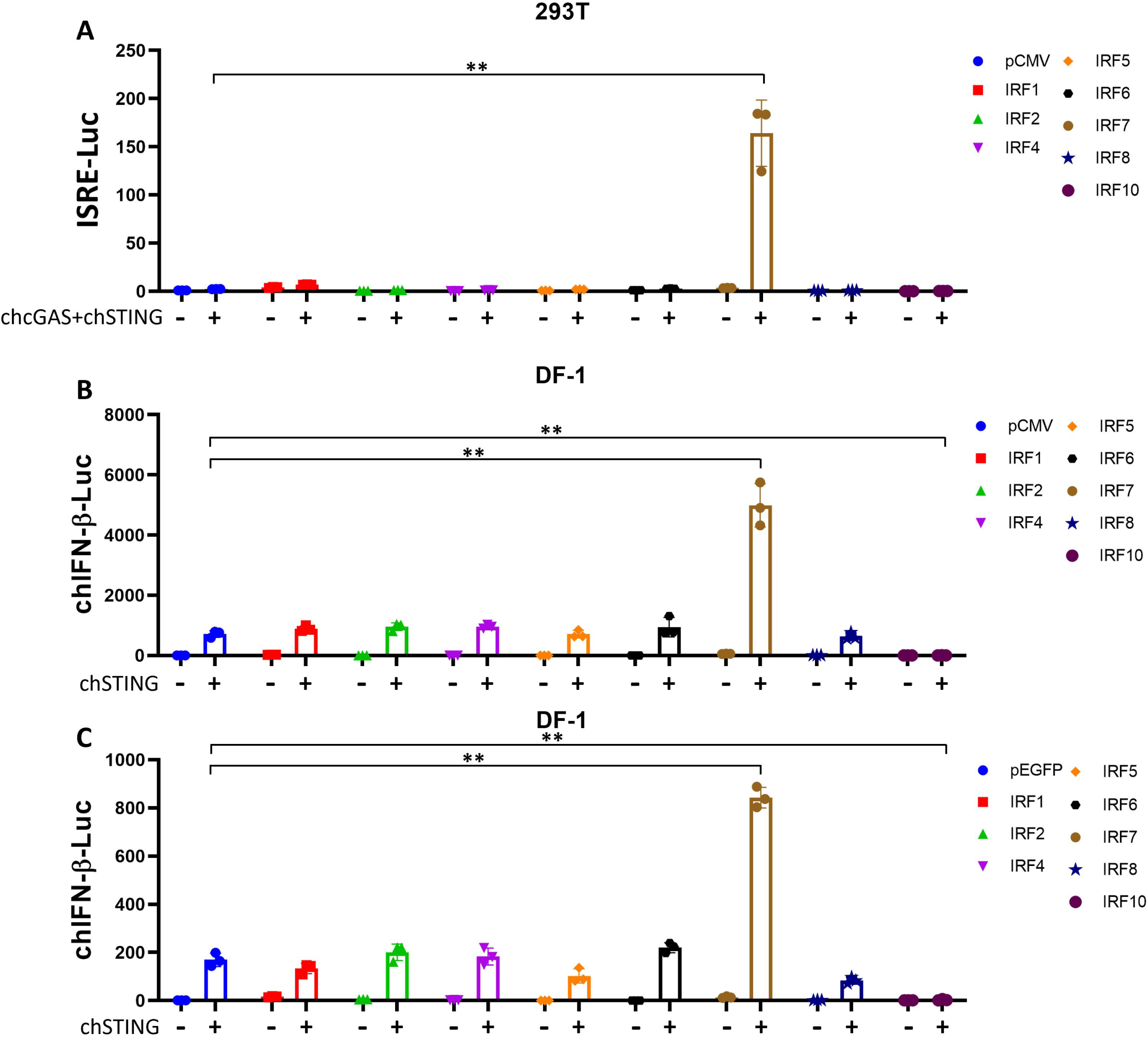
Analysis of chicken (ch) IRF family members in cGAS-STING-IFN signaling by promoter assays. (**A**) Flag-tagged chIRF family members were co-transfected with chGAS-STING into 293T cells, and ISRE promoter activity was measured at 24 h post-transfection. (**B**) Flag-tagged chIRF family members were co-transfected with chSTING into DF-1 cells, and chIFN-β promoter activity was measured at 24 h post-transfection. (**C**) GFP-tagged chIRF family members were co-transfected with chSTING into DF-1 cells, and chIFN-β promoter activity was measured at 24 h post-transfection. ** *p* < 0.01 versus vector controls.

### chIRF10 acts as a negative regulator of the chicken cGAS-STING-IFN signaling pathway

Subsequently, the negative regulatory function of chIRF10 was further characterized. Firstly, we used the chicken macrophage cell line HD11 and treated with 2’3’-cGAMP (an agonist of STING) and poly dA:dT (an agonist of cGAS) to activate cGAS-STING signaling pathway. The results showed that chIRF10 dose-dependently suppresses the activation of the chIFN-β promoter induced by both agonists (Fig 2A and 2B). Similarly, chIRF10 also inhibited chSTING-activated chIFN-β promoter in a dose-dependent manner in DF-1 cells (Fig 2C). Secondly, RT-qPCR results demonstrated that chIRF10 significantly reduces the mRNA expressions of downstream effector genes (IFN-β, MX1, and OASL) activated by either cGAMP or poly dA:dT in HD11 cells (Fig 2D-F). Thirdly, in DF-1 cells, chIRF10 dose-dependently inhibited the mRNA expressions of downstream antiviral genes (IFN-β, MX1, OASL, and PKR) activated by chSTING (Fig 2G-J). Collectively, these results further confirmed that chIRF10 acts as a negative regulator of the chicken cGAS-STING-IFN signaling pathway.

**Figure 2.**
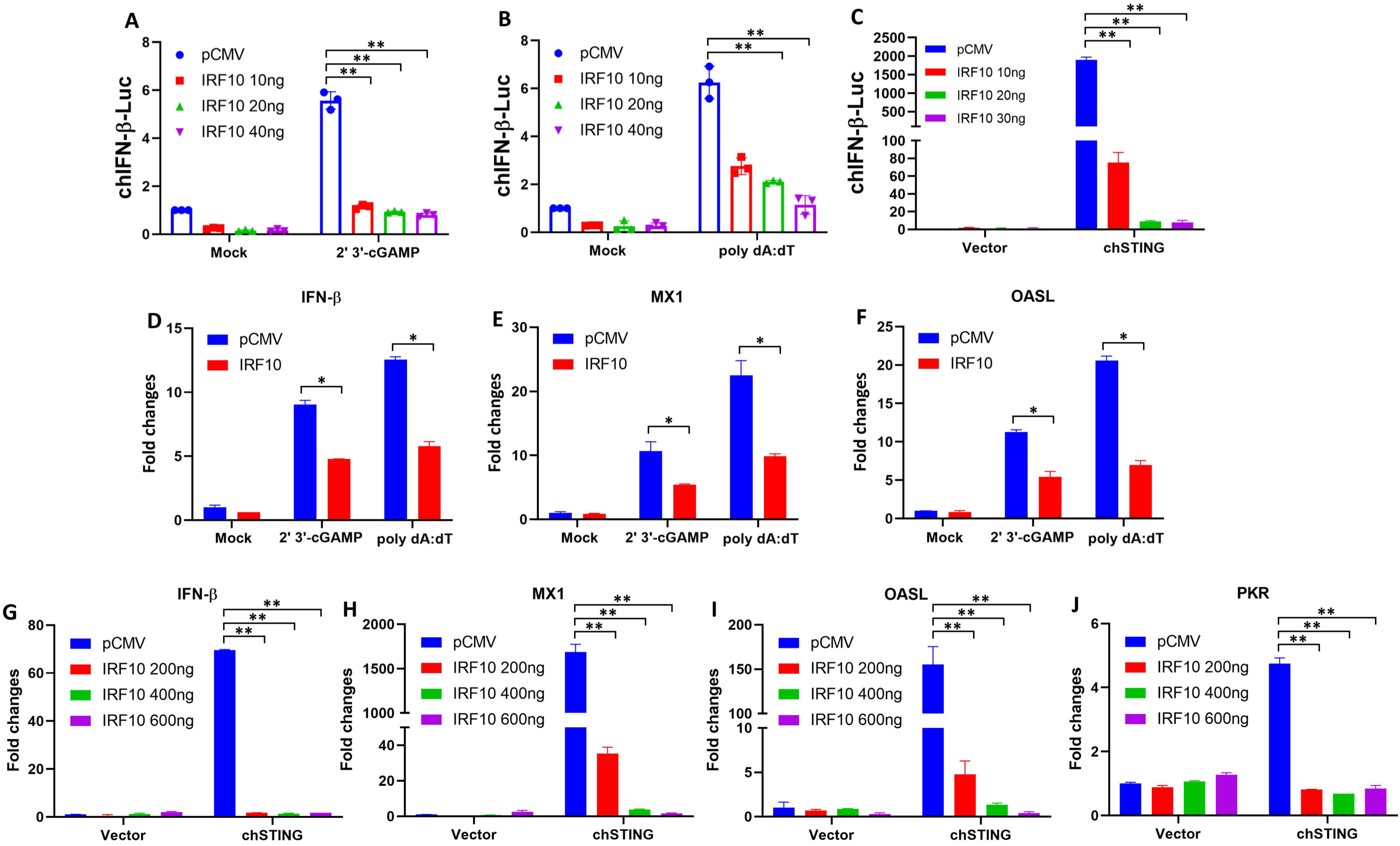
chIRF10 negatively regulates the chicken cGAS-STING-IFN signaling. (**A** and **B**) HD11 cells were transfected with increasing doses of chIRF10 which was normalized with pCMV vector. At 12 h post transfection, the cells were stimulated with 2 μg/mL cGAMP (A) or 1 μg/mL poly dA:dT (B) for 12 h, followed by measurement of chIFN-β promoter activity. (**C**) DF-1 cells were co-transfected with chSTING and increasing doses of chIRF10, normalized with pCMV vector. Cells were collected at 24 h post transfection to measure chIFN-β promoter activity. (**D**-**F**) HD11 cells were transfected with either chIRF10 or pCMV vector. At 12 h post transfection, the cells were stimulated with cGAMP or poly dA:dT for 24 h, and the mRNA expression levels of downstream genes IFN-β (D), MX1 (E), and OASL (F) were detected by RT-qPCR. (**G**-**J**) DF-1 cells were co-transfected with chSTING and increasing doses of chIRF10 for 48 h, the mRNA expression levels of downstream genes IFN-β (G), MX1 (H), OASL (I), and PKR (J) were measured by RT-qPCR. * *p* < 0.05 and ** *p* < 0.01 versus vector controls.

### chIRF10 inhibits the antiviral effect activated by chicken cGAS-STING signaling

Our previous studies have demonstrated that chicken cGAS-STING exerts broad-spectrum antiviral effects (15, 16). To investigate the impact of chIRF10 on the antiviral function of the cGAS-STING pathway, we utilized two RNA viruses, Newcastle disease virus (NDV) and Avian influenza A virus (H1N1), along with two DNA viruses, Vaccinia viruses SMV and VACV. The viral replication levels were determined as follows: NDV was observed for red fluorescence protein (RFP) which it carries by engineering and detected for RFP expression by Western blotting (WB). H1N1 was detected by WB for nucleoprotein (NP) expression. SMV and VACV were measured by quantitative PCR (qPCR) for viral copy numbers. Results from infected HD11 cells showed that chIRF10 significantly inhibits the antiviral effects activated by cGAMP and poly dA:dT against NDV (Fig 3A-D), H1N1 (Fig 3E and 3F), SMV (Fig 3G and 3H), and VACV (Fig 3I and 3J). Similarly, in infected DF-1 cells, chIRF10 also suppressed the chSTING-activated antiviral activity against NDV (Fig 3K and 3L), SMV (Fig 3M), and VACV (Fig 3N). Finally, the mRNA expression level of chIRF10 following cGAMP stimulation and viral infections was measured. The results demonstrated a significant and dose-dependent upregulation of chIRF10 mRNA upon stimulation and various infections (Fig 3O).

**Figure 3.**
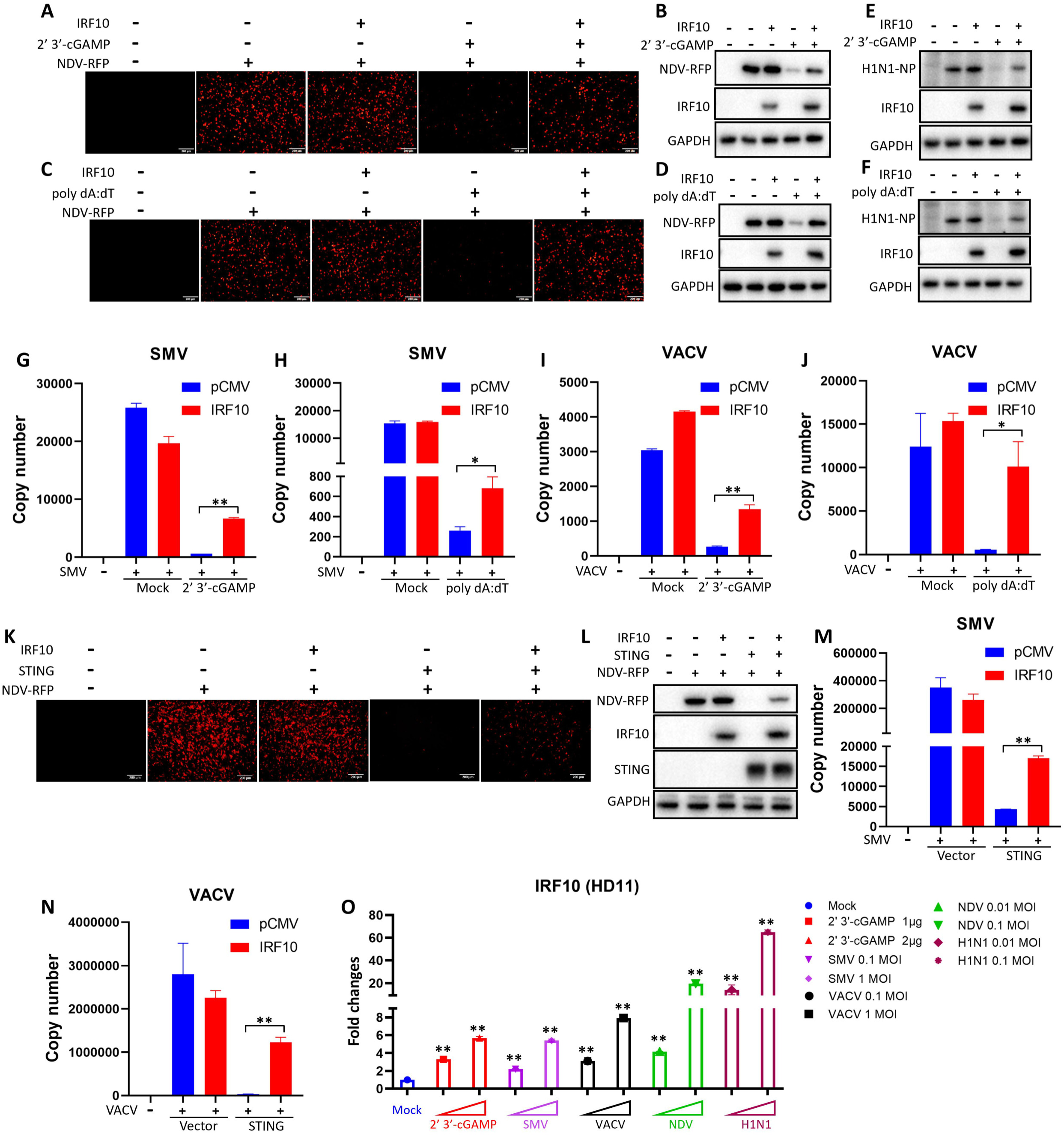
chIRF10 negatively regulates the antiviral function of chcGAS-STING signaling pathway. (**A-D**) HD11 cells were transfected with either chIRF10 or pCMV vector for 12 h, and then stimulated with cGAMP (A and B) or poly dA:dT (C and D) for 12 h, followed by infection with NDV-RFP at 0.01 MOI for 12 h. RFP fluorescence was detected using fluorescence microscopy (A and C), and cell were collected for WB analysis of RFP protein expression (B and D). (**E** and **F**) HD11 cells were transfected and stimulated as in A, followed by infection with AIV (H1N1) at 0.01 MOI for 24 h. The viral NP protein expression was detected by WB. (**G-J**) HD11 cells were transfected and stimulated as in A, followed by infection with SMV at 0.1 MOI for 24 h (G and H) or infection with VACV at 0.1 MOI for 24 h (I and J). The viral copy number was measured by qPCR. (**K** and **L**) DF-1 cells were transfected with chIRF10 and chSTING as indicated for 24 h, and infected with NDV-RFP at 0.01 MOI for 12 h, followed by RFP fluorescence detection by fluorescence microscopy (K) and WB analysis of RFP protein expression (L). (**M** and **N**) DF-1 cells were transfected as indicated for 24 h, and infected with SMV (M) or VACV (N) at 0.1 MOI for 24 h, followed by qPCR measurement of viral copy number. (**O**) HD11 cells were stimulated with cGAMP or infected with the corresponding viruses at the indicated doses. At 24 h post infection, cell were collected and chIRF10 mRNA expression was detected by RT-qPCR. * *p* < 0.05 and ** *p* < 0.01 versus controls.

### The negative regulatory activity of chIRF10 depends on its IRF associated domain

The IRF family members contain two key domains: a DNA-binding domain (DBD) and an IRF associated domain (IAD). The DBD enables IRF to bind promoter DNA, while the IAD facilitates the formation of homodimers or heterodimers with itself or other proteins (43). The DBD of chIRF10 resides in amino acids 6-116, and its IAD is located within amino acids 203-379 (20) (Fig. 4A). Accordingly, chIRF10 ΔDBD and ΔIAD mutants were constructed to investigate their effects on chSTING-activated IFN signaling and antiviral activity. Chicken IFNβ promoter assay results revealed that deletion of the IAD abolished chIRF10’s negative regulatory activity on chSTING-induced IFN signaling, whereas deletion of the DBD did not have such an effect (Fig. 4A). Meanwhile, RT-qPCR results demonstrated that the deletion of the IAD abolished IRF10’s suppressive effect on the expressions of downstream genes IFN-β, MX1 and OASL following chSTING activation, while the deletion of the DBD had no impact (Fig. 4B-D). Finally, antiviral assays demonstrated that chIRF10 ΔIAD had no impact on the antiviral capacity activated by chSTING against NDV (Fig. 4E and 4F), SMV (Fig. 4G), or VACV (Fig. 4H). In contrast, chIRF10 ΔDBD still partially suppressed chSTING mediated antiviral activity, although to a lesser extent (Fig. 4E–H). Collectively, these results suggested that the negative regulatory function of chIRF10 depends on its IAD domain.

**Figure 4.**
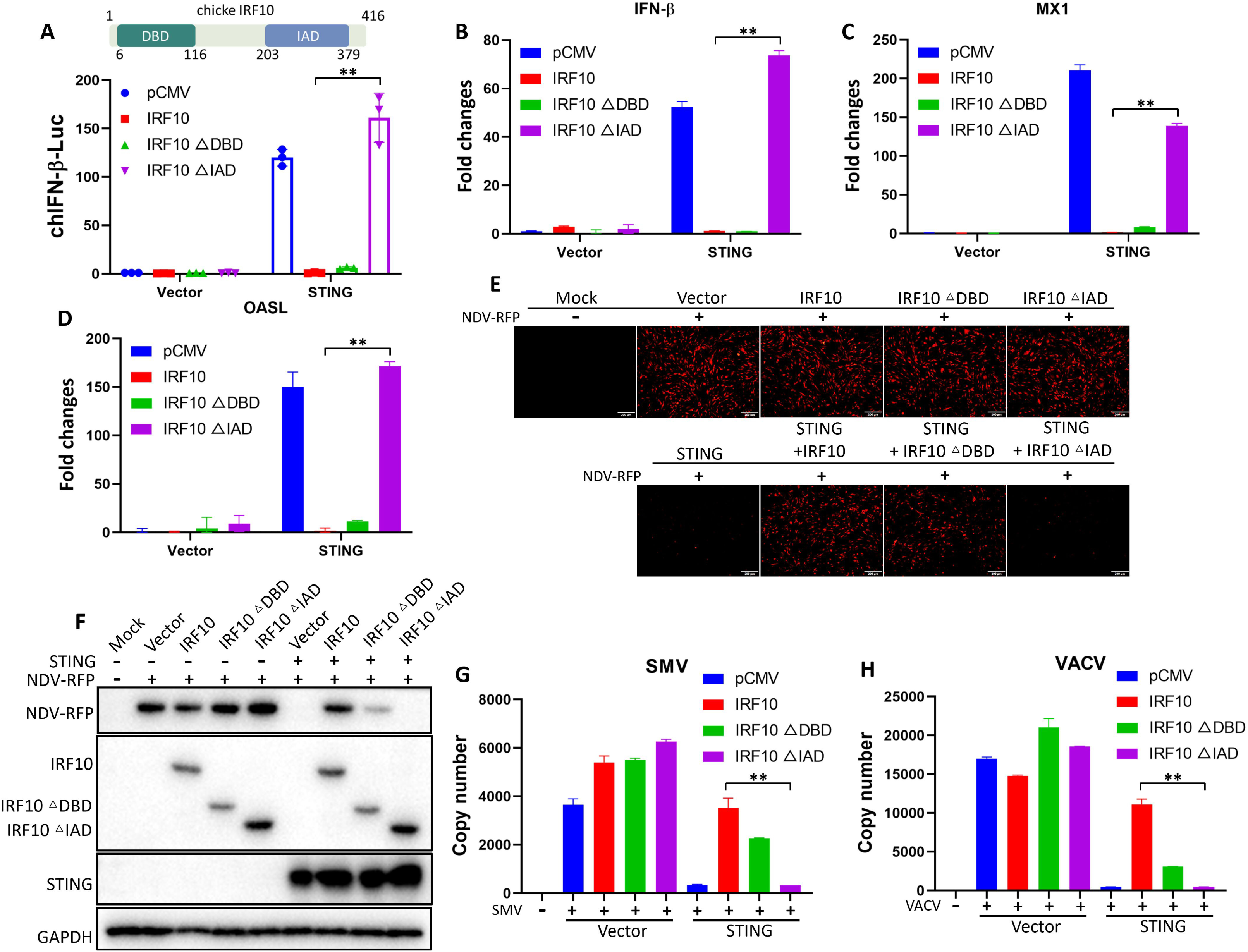
Deletion of the IAD, but not the DBD, abolishes the negative regulatory activity of chIRF10. (**A**) DF-1 cells were co-transfected with chIRF10 or its deletion mutants, with or without chSTING for 24 h, followed by measurement of chIFN-β promoter activity. (**B-D**) DF-1 cells were transfected as in A for 48 h, and the mRNA expressions of chSTING activated downstream genes IFN-β (B), MX1 (C) and OASL (D) were detected by RT-qPCR. (**E** and **F**) DF-1 cells were transfected as indicated for 24 h and infected with NDV at 0.01 MOI for 12 h. RFP fluorescence was observed using fluorescence microscopy (E) and RFP protein expression was detected by WB (F). (**G** and **H**) DF-1 cells were transfected as indicated for 24 h and infected with SMV (G) or VACV (H) at 0.1MOI for 24 h. The viral copy number was measured by qPCR. ** *p* < 0.01 versus controls.

### chIRF10 suppresses IFN signaling activated by downstream mediators of chSTING, specifically chIRF7

TBK1 and IKKε are crucial protein kinases downstream of STING, while chIRF7 is a key transcription factor downstream of chSTING (17, 44). Therefore, expression plasmids for chTBK1 and chIKKε were constructed to examine the effect of chIRF10 on their activation of IFN signaling. The results showed that chIRF10 slightly downregulated the IFN-β promoter activity activated by chTBK1, as well as the mRNA expressions of downstream genes including IFN-β, MX1, OASL, and PKR activated by chTBK1 (Fig 5A). In contrast, chIRF10 significantly inhibited the IFN-β promoter activity and the mRNA expressions of IFN-β, MX1, OASL and PKR activated by chIKKε (Fig 5B). Similarly, chIRF10 significantly and strongly inhibited the IFN-β promoter activity and the mRNA expressions of IFN-β, MX1, OASL and PKR activated by activated by chIRF7 (Fig 5C). These results indicated that the chIRF10 likely targets IRF7 to exert its regulatory role.

**Figure 5.**
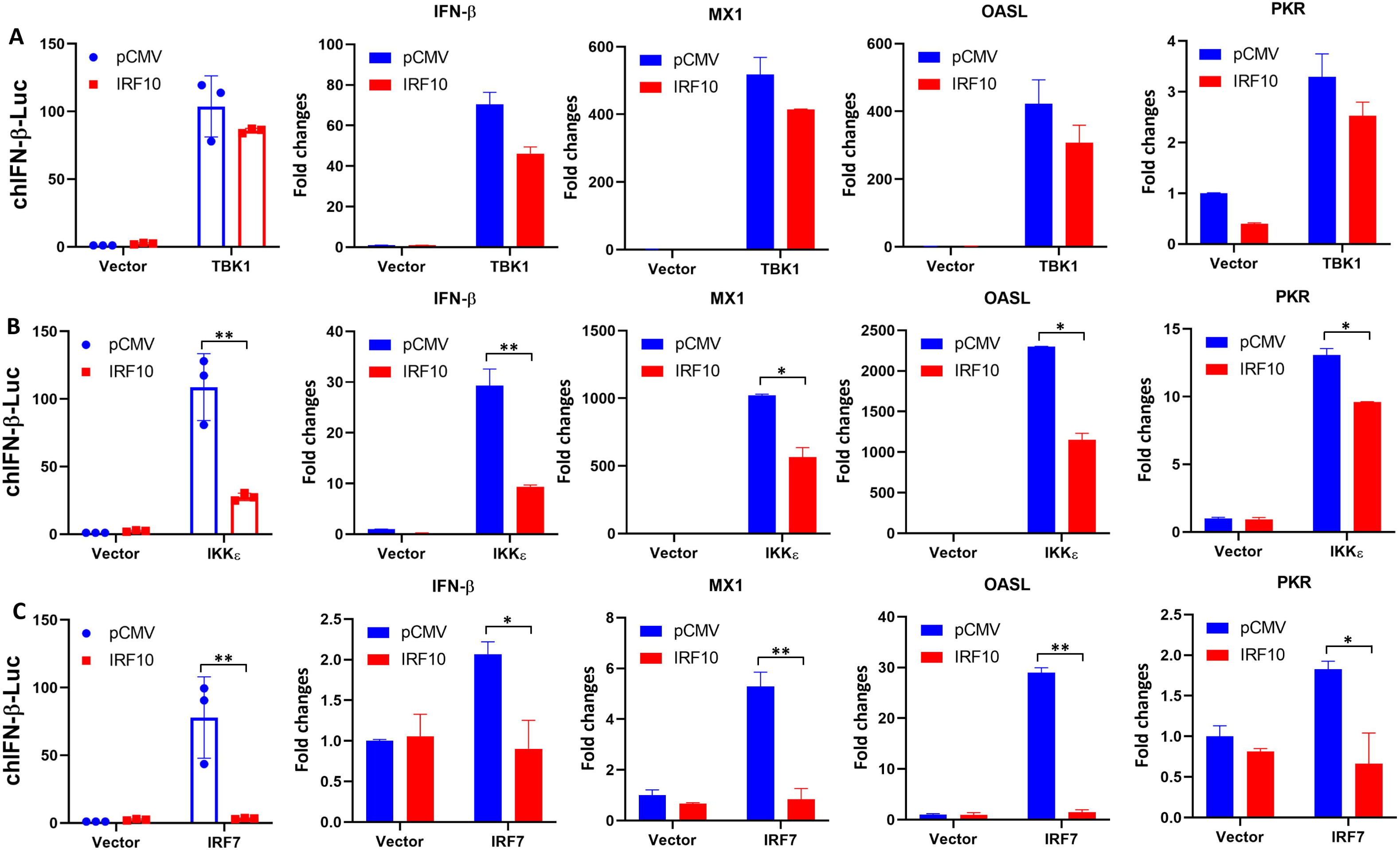
chIRF10 inhibits the IFN signaling activated by chTBK1, chIKKε and chIRF7. (**A**) DF-1 cells were transfected with chIRF10 and chTBK1 as indicated. (**B**) DF-1 cells were transfected with chIRF10 and chIKKε as indicated. (**C**) DF-1 cells were transfected with chIRF10 and chIRF7 as indicated. chIFN-β promoter activity was measured at 24 h post transfection, and the mRNA expression of downstream activated genes IFN-β, MX1, OASL and PKR was detected by RT-qPCR at 48 h post transfection. * *p* < 0.05 and ** *p* < 0.01 versus controls.

### chIRF10 directly targets chIRF7 to suppress its activation

To explore whether chIRF10 targets IRF7 for its negative regulation, we first investigated whether chIRF10 can directly interact with the chIRF7. Confocal microscopy results revealed that both chIRF10 and chIRF10 ΔDBD exhibited significant colocalization with chIRF7, whereas the colocalization was abolished between chIRF10 ΔIAD and chIRF7 (Fig 6A). Similarly, co-immunoprecipitation (Co-IP) assays demonstrated that chIRF10 and chIRF10 ΔDBD, but not chIRF10 ΔIAD, could strongly interact with chIRF7 (Fig 6B and C). Additionally, chIRF10 was also found to interact with chSTING, chTBK1, and chIKKε (Fig. S1).

**Figure 6.**
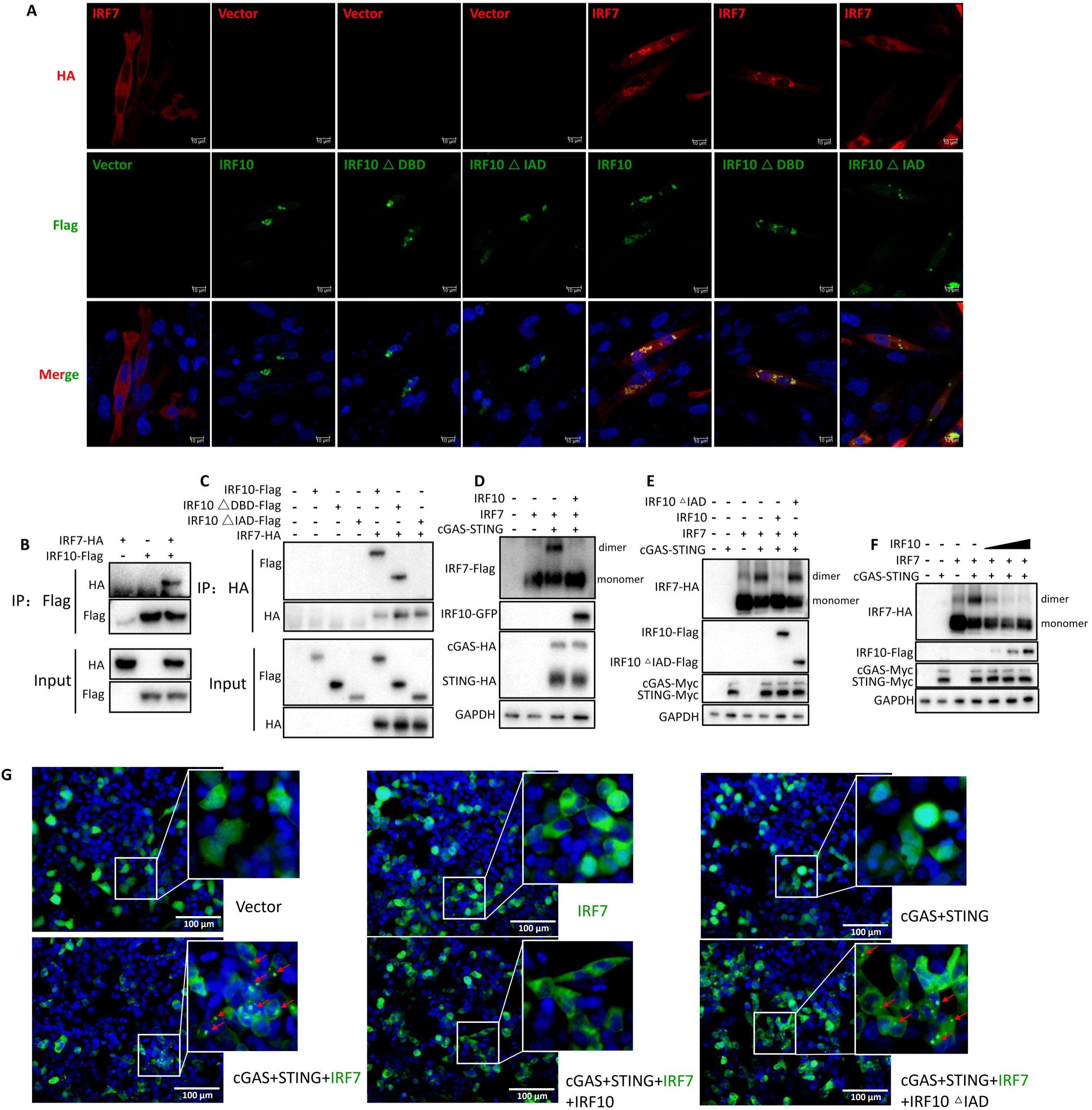
chIRF10 targets chIRF7 and inhibits chIRF7 activation. (**A**) DF-1 cells were transfected with chIRF10 and its deletion mutants, with or without chIRF7 as indicated for 24 h. The co-localization of chIRF10/mutants and chIRF7 was examined by laser scanning confocal microscopy following IFA staining. (**B-C**) 293T cells were transfected with chIRF10 and chIRF7 (B), chIRF10/ chIRF10ΔDBD/chIRF10ΔIAD and chIRF7 (B) as indicated for 48 h. The protein interactions between chIRF10/mutants and chIRF7 were assessed by Co-IP using the indicated antibodies. (**D-F**) 293T cells were transfected with chIRF10, chIRF7 plus chcGAS-STING (D), chIRF10/chIRF10ΔIAD, chIRF7 plus chcGAS-STING (E), increasing doses of chIRF10, chIRF7 plus chcGAS-STING (F) as indicated for 24 h. The chIRF7 dimerization levels were analyzed by native PAGE. (**G**) 293T cells were co-transfected with GFP-tagged chIRF7, Flag-tagged chIRF10/chIRF10ΔIAD, HA-tagged chcGAS-STING as indicated. The puncta formation of chIRF7 indicating its activation was observed by fluorescence microscopy 24 h post transfection, and marked as red arrows.

Next, we examined whether chIRF10 inhibits the activation of chIRF7. The results showed that chIRF10 markedly suppressed chcGAS-STING induced dimerization of chIRF7 (Fig 6D), and this inhibition was dependent on the IAD domain of chIRF10 (Fig 6E). Further validation indicated that chIRF10 inhibited chIRF7 dimerization in a concentration-dependent manner (Fig 6F). Interestingly, in 293T cells, co-transfection of chcGAS-chSTING with chIRF7 promoted the formation of distinct speckles by chIRF7, which was abolished upon the addition of chIRF10 but reappeared when chIRF10 ΔIAD was introduced (Fig 6G). These speckles likely represent dimerized or oligomerized forms of chIRF7, indicative of its activation. In summary, consistent with the signaling activities of DBD and IAD deletion mutants of chIRF10, the results clearly demonstrated that chIRF10 interacts with chIRF7 via its IAD domain and inhibits IRF7 activation.

### chIRF10 knockout enhances the cGAMP-triggered signaling activation and suppresses viral replication in HD11 cells

To further elucidate the negative regulation of chIRF10, it was knocked out in HD11 cells and IRF10^-/-^ cells were obtained. Upon cGAMP stimulation, the absence of chIRF10 was found to enhance the mRNA expressions of cGAMP activated IFN-β, OASL, and MX1 (Fig. 7A-C). Secondly, two obtained chIRF10^-/-^ HD11 cell clones were infected with NDV, revealing a significant reduction in viral replication in both clones (Fig. 7D-G). Thirdly, the replication levels of SIV (Fig. 7H), SMV (Fig. 7I), and VACV (Fig. 7J) were also markedly decreased in the chIRF10^-/-^ HD11 cells. These findings further established that endogenous chIRF10 acts as a negative regulator of the cGAS-STING-IFN antiviral axis.

**Figure 7.**
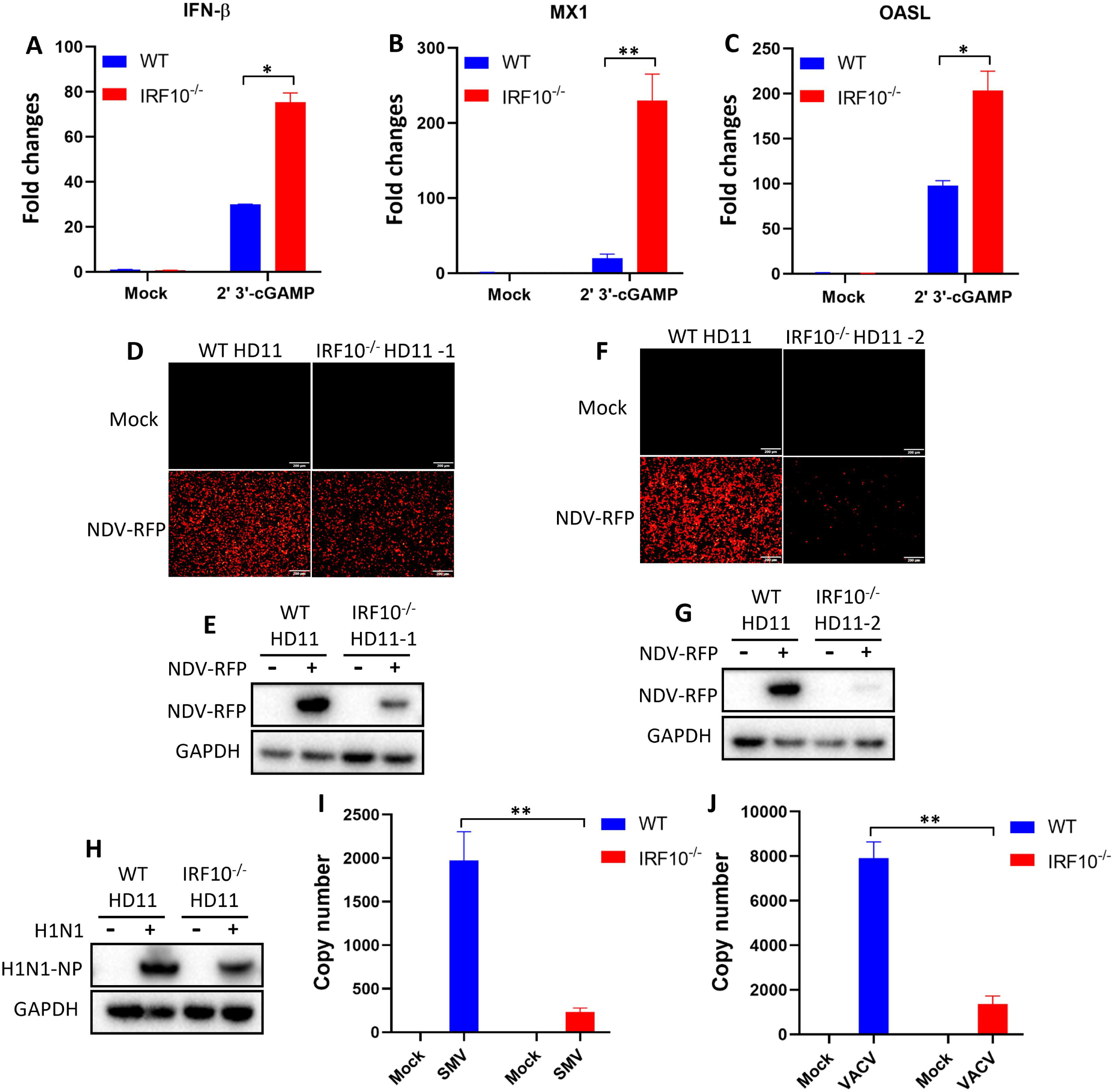
The chcGAS-STING antiviral signaling is upregulated in chIRF10^-/-^ HD11 cells. (**A-C**) Wild-type (WT) and chIRF10^-/-^ HD11 cells were stimulated with cGAMP for 24 h, and the mRNA expression levels of activated IFN-β (A), MX1 (B), and OASL (C) were detected by RT-qPCR. (**D-G**) Two independent clones of chIRF10^-/-^ HD11 cells plus WT HD11 cells were infected with NDV at 0.01 MOI for 12 h. RFP fluorescence was observed under a fluorescence microscope (D and F), and RFP protein expression was detected by WB (E and G). (**H-J**) chIRF10^-/-^ HD11 plus WT HD11 cells were infected with AIV at 0.01 MOI (H), SMV at 0.1 MOI (I), or VACV at 0.1 MOI (J). At 24 h post infection, AIV NP protein expression was detected by WB, and viral copy numbers of SMV and VACV were quantified by qPCR. * *p* < 0.05, ** *p* < 0.01 and ns versus controls.

### Reconstitution of chIRF10 and its mutants in chIRF10^-/-^ HD11 cells re-confirms the negative regulation of chIRF10

It has been established that in the absence of chIRF10, cGAMP-activated IFN signaling is upregulated and viral replication is reduced. To further validate the regulation of chIRF10, chIRF10 and its mutants were reintroduced into chIRF10^-/-^ HD11 cells. First, we examined the cGAMP activated IFN signaling after reconstitution with chIRF10, chIRF10 ΔDBD, and chIRF10 ΔIAD. Compared to wild-type (WT) HD11 cells, the mRNA levels of IFN related genes (IFN-β, OASL, and MX1) were elevated in chIRF10^-/-^ HD11 cells upon cGAMP stimulation (Fig. 8A-C). Notably, reconstitution with chIRF10 and chIRF10 ΔDBD, but not chIRF10 ΔIAD, reduced the expression of these cGAMP induced IFN related genes (Fig. 8A-C). Second, viral replications were assessed after complementation with chIRF10 and its mutants. Relative to WT HD11, chIRF10^-/-^ HD11 cells showed decreased replications of NDV (Fig. 8D and 8E) and AIV (Fig. 8F), and all these viral replications were elevated upon reconstitution with chIRF10 or chIRF10 ΔDBD, but not with chIRF10 ΔIAD (Fig. 8D-F). Consistent results were observed for the replication levels of SMV (Fig. 8G) and VACV (Fig. 8H) after reconstitution. These results further clarified the negative regulatory impact of chIRF10 on chicken cGAS-STING-IFN antiviral signaling pathway.

**Figure 8.**
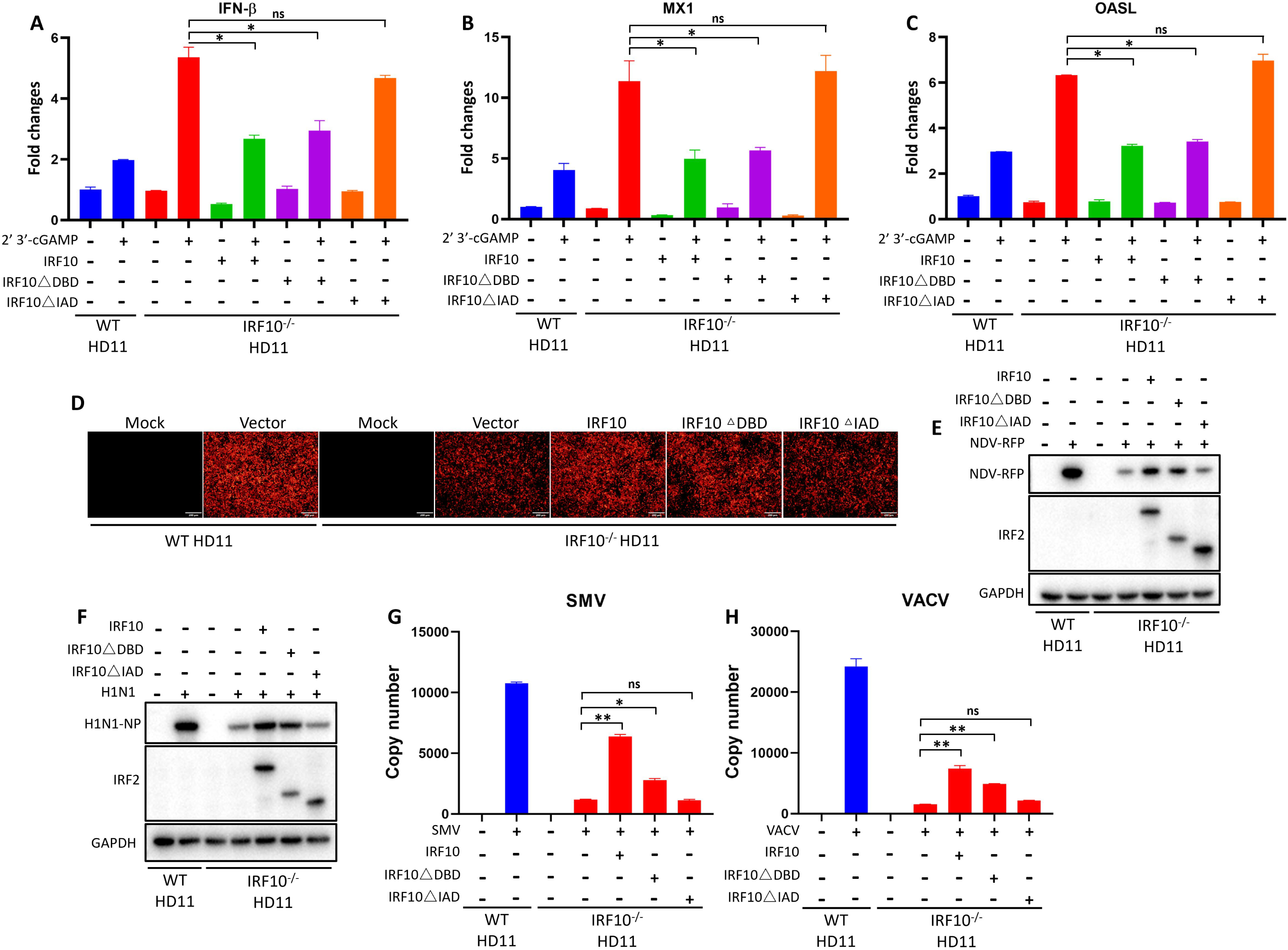
Assessment of the cGAS-STING antiviral signaling in chIRF10 reconstituted IRF10^-/-^ HD11 cells. (**A-C**) WT and reconstituted chIRF10^-/-^ HD11 cells transfected with chIRF10, chIRF10 ΔDBD, chIRF10 ΔIAD or empty vector for 12 h, were stimulated with cGAMP for 24 h. Cells were harvested and the mRNA expression levels of IFN-β (A), MX1 (B), and OASL (C) were detected by RT-qPCR. (**D** and **E**) WT and reconstituted chIRF10^-/-^ HD11 cells transfected with chIRF10, chIRF10ΔDBD, chIRF10ΔIAD or empty vector for 12 h, were infected with NDV at 0.01 MOI for 12 h. RFP fluorescence was observed by fluorescence microscopy (D), and RFP protein expression was detected by WB (E). (**F-H**) WT and the constituted chIRF10^-/-^ HD11 cells, as indicated, were infected with AIV at 0.01 MOI (F), SMV at 0.1 MOI (G), or VACV at 0.1 MOI (H) for 24 h. The NP protein expression of AIV was detected by WB (F), and viral copy numbers of SMV (G) and VACV (H) were quantified by qPCR. * *p* < 0.05, ** *p* < 0.01 and ns versus controls.

## Discussion

It has been extensively demonstrated that chIRF7 acts as the primary IRF downstream of chSTING, responsible for binding to the IFN-β promoter (17, 18). However, whether other chicken IRF family members, besides chIRF7, are involved in the chicken cGAS-STING-IFN antiviral signaling pathway remains unknown. Given the limited research on chicken IRFs, this study first employed a promoter screening assay and revealed that only chIRF7 mediates IFN signaling triggered by chicken cGAS-STING, which is consistent with previous findings (Fig. 1A) (17, 18). Additionally, we reported for the first time that chIRF10, which is unique to chickens, significantly negatively regulates the chicken cGAS-STING-IFN signaling pathway (Fig. 1B and 1C).

Although chIRF10 was identified as early as 2002, its functional characterization has remained scarce (20, 45). Here, we unveil for the first time the negative regulatory role of chIRF10 in the chicken cGAS-STING-IFN antiviral response and further elucidate its underlying mechanism. chIRF10 potently suppresses cGAS-STING activated IFN signaling in two chicken cell lines (DF-1 and HD11) (Fig. 2). Our previous studies have reported the broad-spectrum antiviral activity of chicken cGAS-STING, which is further corroborated in this study (Fig. 3) (15, 16). Further investigation into the function of chIRF10 demonstrated that it significantly inhibits the antiviral signaling activated by chicken cGAS-STING (Fig. 3). Moreover, this study found that chIRF10 expression can be induced by viral infection (Fig. 3O), suggesting that chIRF10 may function as an IFN-stimulated gene (ISG) and could potentially be exploited by viruses to suppress innate immune responses and facilitate immune evasion. Unfortunately, due to the lack of specific antibodies, the expression level of chIRF10 following viral infection could not be detected at the protein level.

Similar to other IRF family members, chIRF10 contains a DNA binding domain (DBD, 6-116 aa) and an IRF associated domain (IAD, 203-379 aa) (Fig. 4A). This study found that it is the deletion of the IAD, rather than the DBD, that abolishes the inhibitory effect of chIRF10 on chSTING mediated IFN activation (Fig. 4A-D). Furthermore, the loss of the IAD also eliminates the negative regulatory effect of chIRF10 on the antiviral response triggered by chSTING (Fig. 4E-H).

chTBK1 and chIRF7 are key kinase and transcription factor, respectively, downstream of chSTING, and IKKε has also been identified as an important kinase in the mammalian STING pathway (17, 44, 46). This study demonstrates that chIRF10 exerts varying degrees of inhibition on IFN signaling activated by downstream components of chSTING, including chTBK1 (Fig. 5A), chIKKε (Fig. 5B), and chIRF7 (Fig. 5C). Intriguingly, while chIRF10 strongly inhibits chIRF7, it exhibits only weak inhibition toward chTBK1, suggesting the potential existence of additional transcription factors downstream of chTBK1 responsible for the transcriptional activation of type I IFN.

It is known that the IAD of IRFs mediates interactions with other proteins (43). In our study, chIRF10 was found to significantly and strongly inhibit IFN signaling activated by chIRF7, and this regulatory activity is dependent on its IAD (Fig 4 and 5). Therefore, we investigated the interaction between chIRF10 (and its mutants) and chIRF7. We found that chIRF10 strongly interacts with chIRF7 in a manner dependent on the IAD (Fig. 6A-C). Similar to IRF3, IRF7 forms dimers that translocate into the nucleus to initiate type I IFN transcription. This study demonstrates that chIRF10 inhibits the formation of chIRF7 dimers induced by chicken cGAS-STING, and this inhibition also requires the IAD (Fig. 6D-F). Interestingly, we observed that chicken cGAS-STING expression in 293T cells promotes the formation of distinct punctate structures by GFP-tagged chIRF7 (Fig. 6G). We thought that this punctate pattern represents the oligomeric state of chIRF7, indicative of its activation. Furthermore, chIRF10, but not chIRF10 ΔIAD, suppresses this punctate formation, providing additional evidence at the cellular level for the negative regulatory role of chIRF10 toward chIRF7 (Fig. 6G).

Finally, knockout of chIRF10 in HD11 cells resulted in significantly enhanced IFN signaling activated by cGAMP, along with markedly reduced viral replication levels in the knockout cells (Fig. 7). Following the complementation of chIRF10^-/-^ HD11 cells with wild-type chIRF10 or the chIRF10 ΔDBD, a significant suppression of cGAMP-induced IFN signaling and a concomitant increase in viral replications were observed. In contrast, cells complemented with chIRF10 ΔIAD exhibited no such effects. (Fig. 8). These results further confirm the negative regulatory role of chIRF10 in the cGAS-STING-IFN antiviral signaling pathway.

In summary, we are the first to identify the negative regulatory function of chIRF10 in the cGAS-STING-IFN antiviral pathway and elucidate the underlying mechanism, in which the chIRF10 targets chIRF7 and thereby suppresses its activation (Fig. 9). Our study expands the understanding of the IRF family, particularly chicken IRF members, deepens the knowledge of the cellular function of chIRF10, fills a gap in the regulation of antiviral responses in chickens, and provides a theoretical basis for the prevention and control of viral diseases in poultry.

**Figure 9.**
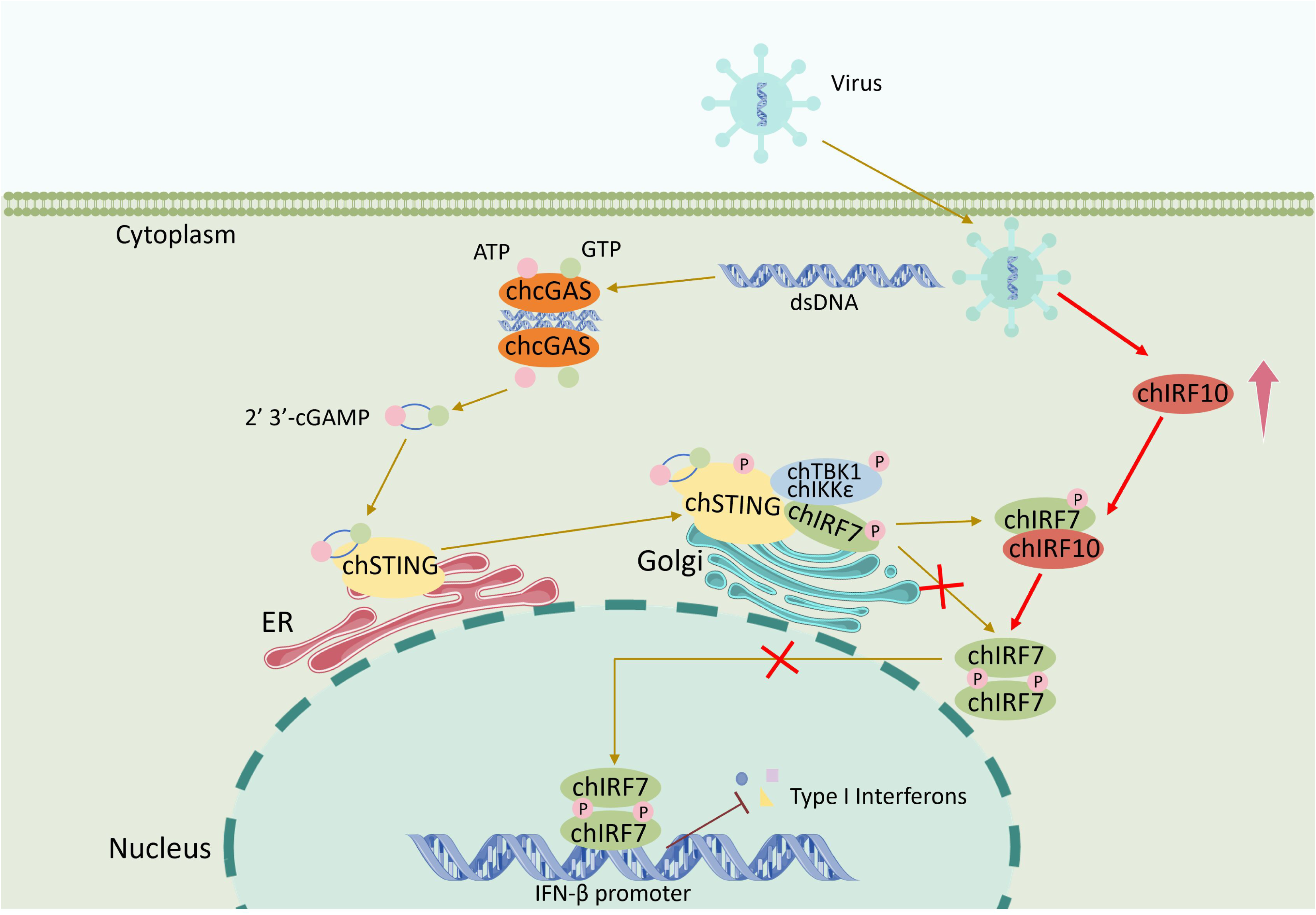
Schematic mechanism of action for the chIRF10 negative regulation of chicken cGAS-STING-IFN antiviral signaling. The invading virus directly releases its genomic double-stranded DNA (dsDNA) into the cytosol, or damages host cells causing the release of mitochondria or nuclear DNA into the cytosol, where the DNA is recognized by the DNA receptor chçGAS. Upon binding dsDNA, chçGAS catalyzes ATP and GTP into the 2’3’-cGAMP, which, as a second messenger, activates chSTING on the endoplasmic reticulum (ER). Activated chSTING translocates from ER to Golgi apparatus, allowing chSTING to recruit and activate the kinases chTBK1/IKKε and transcription factor chIRF7. The phosphorylated chIRF7 forms a dimer, translocates from the cytoplasm into the nucleus, and binds the type I IFN promoter to initiate IFN transcription. In contrast, during viral infection, the chIRF10 is upregulated. chIRF10 interacts with chIRF7 via its IAD domain, and inhibits chIRF7 activation, thereby negatively regulating the chicken cGAS-STING-IFN antiviral signaling.

## Materials and methods

### Antibodies and reagents

The FLAG Rabbit mAb (14793) and HA Rabbit mAb (3724) were acquired from Cell Signaling Technology (Boston, MA, USA). The mouse anti-FLAG mAb, mouse anti-HA mAb, mouse anti-GAPDH mAb and mouse anti-GFP mAb were all acquired from Transgen Biotech (Beijing, China). Mouse anti-Myc mAb (60003-2-Ig) was purchased from ProteinTech (Wuhan, China). Anti-RFP mAb HRP-DirecT (M204-7) was purchased from MBL Beijing Biotech (Beijing, China). HRP goat anti-rabbit IgG (H+L) highly cross-adsorbed secondary antibody and goat anti-mouse IgG (H+L) highly cross-adsorbed secondary antibody were all obtained from Sangon Biotech (Shanghai, China). Goat anti-mouse IgG H&L Alexa Fluor®594 (ab150120) were acquired from Abcam (Cambridge, UK). Goat anti-rabbit IgG (H+L) cross-adsorbed 488 (35553) were from ThermoFisher Scientific (Shanghai, China).

Lipofectamine 2000 were purchased from Thermo Fisher (Sunnyvale, CA, USA). TransIT®-LT1 Transfection Reagent (MIR 2300) for HD11 was purchased from Mirus Bio (Madison, WI, USA). TRIpure Reagent was bought from Aidlab (Beijing, China). Double-luciferase reporter assay kits, 2 × Phanta Max Master Mix(P515-01), HiScript® 1st Strand cDNA Synthesis Kit, ChamQ Universal SYBR qPCR Master Mix and 2×Taq Master Mix (Dye plus) were all from Vazyme Biotech (Nanjing, China). The 2×MultiF Seamless Assembly Mix was acquired from Abclonal (Wuhan, China). 2’3’-cGAMP and polydA:dT were purchased from InvivoGen (Hong Kong, China). Protein A/G PLUS-Agarose was bought from Santa Cruz Biotechnology (sc-2003, CA, USA). 4’,6-diamidino-2-phenylindole (DAPI) staining solution (C1005) was from Beyotime (Shanghai, China).

### Cell transfection and viruses

The human embryo kidney HEK-293T cells (ATCC Cat# CRL-3216), and chicken fibroblast DF-1 cells (ATCC Cat# CRL-12203) were maintained in DMEM (Hyclone Laboratories, USA) supplemented with 10% fetal bovine serum (FBS, Vazyme Biotech) and 1% penicillin/streptomycin, and maintained at 37 °C with 5% CO2. The chicken macrophages HD11 cells (BCRJ-Linhagens celulares Cat# 0099) was cultured in RPMI 1640 medium (Hyclone Laboratories) supplemented with 10% FBS and 1% penicillin/streptomycin, and maintained at 37 °C with 5% CO2. Transfection was performed using the Lipofectamine 2000 or TransIT-LT1 Transfection Reagent following the manufacturer’s instructions. Newcastle disease virus (NDV)-RFP, avian influenza virus (AIV, H1N1) and vaccinia viruses (strains VACV and SMV) were all preserved in our laboratory.

### Gene cloning and gene mutations

The chicken (ch) cGAS, chSTING, chIRF7, and chIRF1 plasmids were previously constructed and have been used in our laboratory. Total RNA was extracted from HD11 cells using TRIpure reagent, and cDNA was prepared by reverse transcription from total RNA. The chIRF2 (NM_205196), chIRF4 (NM_204299), chIRF5 (NM_001031587), chIRF6 (XM_025143814), chIRF8 (NM_205416), chIRF10 (NM_204558), chTBK1 (NM_001199558), chIKKε (XM_428036) open reading frames (ORFs) were amplified by PCR from prepared cDNA using the designed primers as shown in Supplementary Table 1. The PCR products of chIRF were cloned into the *Bgl* II and *Kpn* I sites of pEGFP-C1 vector and the *EcoR* I and *Sal* I sites of p3×FLAG-CMV-7.1 vector, respectively, by seamless cloning. The PCR products of chTBK1 and chIKKε were cloned into the *EcoR* I and *EcoR V* sites of pCAGGS-2HA vector by seemless cloning. The mutation PCR primers of chIRF10 were designed by QuickChange Primer Design method (https://www.agilent.com) which were shown in Supplementary Table 2. The mutation PCR was performed with Phanta Max Master Mix and p3×FLAG-CMV-chIRF10 as the template, as we described previously (47).

### Promoter luciferase reporter gene assay

293T or DF-1 cells grown in 96-well plates (2-3×10^4^ cells/well) were co-transfected using Lipofectamine 2000 with ISRE-luciferase or chicken IFN-β-luciferase reporter (10 ng/well) and β-actin Renilla luciferase (Rluc) reporter (0.2 ng/well), together with the indicated plasmids or vector control (5–50 ng/well). HD11 cells were similarly transfected with chicken IFN-β Fluc using TransIT-LT1 transfection reagent. The total DNA per well was normalized to 50 or 60 ng by adding corresponding empty vectors. With or without further stimulation post transfection, the cells were lysed, and the Fluc and Rluc activities were measured successively by double-luciferase reporter assay kit, as we described previously (47).

### DNA and RNA extraction, reverse transcription and quantitative PCR (qPCR)

Total cellular DNA was extracted using the HiPure Tissue DNA Mini kit (Magen, Guangzhou, China), and total RNA was extracted using TRIpure reagent following the manufacturer’s recommendations. The extracted RNA was reverse transcribed into cDNA by using HiScript 1st strand cDNA synthesis kit. The DNA and cDNA were used for measuring target gene expressions by quantitative PCR with SYBR qPCR master Mix (Vazyme, Nanjing, China) using StepOne Plus equipment (Applied Biosystems). The qPCR program contained a denaturation step at 95°C for 30 s, and 40 cycles of 95°C for 5 s and 60 °C for 30 s. The relative mRNA levels were calculated using 2^−ΔΔCT^ method after normalization to GAPDH mRNA levels. The sequences of qPCR primers were shown in Supplementary Table 3.

### Western blotting (WB) and co-immunoprecipitation (Co-IP) analysis

The protein lysate samples from RIPA extraction were mixed with 4×loading buffer at a ratio of 3:1 and boiled for 5-10 min. The supernatants after centrifugation were run by SDS-PAGE, and then the proteins were transferred to PVDF membrane. The membrane was incubated with 5% skim milk solution at room temperature (RT) for 2 h, probed with the indicated primary antibodies at 4℃ overnight, and then incubated with secondary antibodies for 1 h at RT, followed by detection with ECL substrate and imaging system (Tanon, Shanghai, China).

For co-immunoprecipitation (Co-IP), the cleared cell lysates from transfected cells in 6-well plate (6-8×10^5^ cells/well) were incubated with 1 μg of specific antibody at 4°C overnight with shaking, followed by incubation with Protein A/G PLUS-Agarose for 2-3 h. The agarose was washed and eluted with 40 μL of 2 × SDS sample buffer. The elution samples together with input lysate inputs were both subjected to Western blotting.

### Native polyacrylamide gel electrophoresis and chIRF7 dimerization assay

293T cells plated in 12-well plate (3-4×10^5^ cells/well) were transfected with the indicated combinations of chcGAS, chSTING, chIRF7, chIRF10 and chIRF10 △ IAD for 24 h. Cells were lysed in non-denaturing lysis buffer for 15 min, the protein supernatant was collected and mixed with 2× non-denaturing loading buffer. Meantime, the non-denaturing polyacrylamide gel (without SDS) was pre-electrophoresed at 100 V for 30 min until the current stabilized, with the inner electrophoresis tank buffer containing 1% sodium deoxycholate. Next, the non-denatured protein samples were then loaded and electrophoresed at 120 V for 60 minutes on an ice bath. Subsequent steps followed the standard Western blotting procedures.

### Confocal microscopy

DF-1 cells grown on 15 mm glass bottom cell culture dish (5×10^5^ cells) were co-transfected with chIRF7-HA or pCAGGS-2HA along with either chIRF10-Flag, chIRF10 ΔDBD-Flag, chIRF10 ΔIAD-Flag or p3×FLAG-CMV, using Lipofectamine 2000. At 24 h post transfection, the cells were fixed with 4% paraformaldehyde at RT for 30 min, and permeabilized with 0.5% Triton X-100 for 20 min. After washing with PBS, the cells were sequentially incubated with Flag rabbit mAb (1:200) and HA mouse mAb (1:200), and next secondary Goat anti-rabbit IgG (H+L) cross-adsorbed 488 (1:800), Goat anti-mouse IgG H&L Alexa Fluor 594 (1:800). The stained cells were counter-stained for cell nucleus with 0.5mg/mL 4’,6-diamidino-2-phenylindole (DAPI) at 37°C for 15 min. Lastly, cells were visualized under laser-scanning confocal microscope (LSCM, Leica SP8, Solms, Germany) at the excitation wavelengths 488 nm and 594 nm, respectively.

### Gene knockout (KO) macrophage cell clones made by CRISPR gRNAs

The CRISPR gRNAs targeting chIRF10 were designed using the web tool from Benchling (www.benchling.com). Two pairs of gRNAs were selected according to the prediction score, with the sequences of gRNA encoding sequences are shown in Supplementary Table 4. The annealed gRNA encoding DNA sequences were inserted into the *Bbs*I site of pX458-EGFP and the resultant pX458-gRNA plasmids were DNA sequence confirmed. HD11 cells grown in 6-well plates (6-8×10^5^ cells/well) were transfected with pX458-gRNA using TransIT-LT1, respectively. At 24 h post transfection, flow cytometric sorting was performed and the sorted GFP positive cells were grown in 96-well plates for subsequent monoclonal growth by limiting dilution. The single clones of HD11 cells were detected for genomic DNA editing by PCR using the primers shown in Supplementary Table 4. If necessary, the PCR products were run by non-denatured polyacrylamide gel electrophoresis following by silver staining. The genomic PCR products were cloned, sequenced, and aligned to analyze the base insertion and deletion (ins/del) mutations, as we described previously (47). Finally, homologous IRF10 KO (IRF10^-/-^) HD11 cell clones were obtained.

### Statistical analysis of experimental data

All the results represent more than two similar experiments. The data of bar graphs were showed as the mean ± standard deviation (SD) and analyzed by Student’s *t*-test and *ANOVA*, as propriate, for statistical analysis. Statistical signs were as follows: * *p* < 0.05, ** *p* < 0.01 and ns. The *p* < 0.05 was considered statistically significant, and ns denotes no statistically significance.

## Author contribution statement

S.J and J.Z conceived and designed the experiments; S.J, J.H, K.W, M.L, H.H and B.S performed the experiments; N.C, W.Z, and J.Z analyzed the data; S.J and J.Z wrote the paper. All authors have contributed to the manuscript and approved the submitted version.

## Data Availability Statement

The WB raw data, IFA raw data and the numerical raw data of this study are all available upon request.

